# A chromosome level genome assembly of longnose gar, *Lepisosteus osseus*

**DOI:** 10.1101/2022.12.21.521478

**Authors:** Rittika Mallik, Kara B. Carlson, Dustin J. Wcisel, Michael Fisk, Jeffrey A. Yoder, Alex Dornburg

## Abstract

Holosteans (gars and bowfins) represent the sister lineage to teleost fishes, the latter being a clade that comprises over half of all living vertebrates and includes important models for comparative genomics and human health. A major distinction between the evolutionary history of teleosts and holosteans is that all teleosts experienced a genome duplication event in their early evolutionary history. As holostean genomes did not undergo a round of genome duplication, they have been heralded as a means to bridge teleost models to other vertebrate genomes. However, only three species of holosteans have been genome sequenced to date and sequencing of more species is needed to fill sequence sampling gaps and provide a broader comparative basis for understanding holostean genome evolution. Here we report the first high quality reference genome assembly and annotation of the longnose gar (*Lepisosteus osseus*). Our final assembly consists of 22,709 scaffolds with a total length of 945 bp with contig N_50_ of 116.6 kb. Using BRAKER2, we annotated a total of 30,068 genes. Analysis of the repetitive regions of the genome reveals the genome to contain 29.1% transposable elements, and the longnose gar to be the only other known vertebrate outside of the spotted gar to contain CR1, L2, Rex1, and Babar. These results highlight the potential utility of holostean genomes for understanding the evolution of vertebrate repetitive elements and provide a critical reference for comparative genomic studies utilizing ray-finned fish models.

**Significance:** Over half of all living vertebrates are teleost fishes, including numerous experimental models such as zebrafish (*Danio rerio*) and medaka (*Oryzias latipes*). However, translating research in teleost models to other organisms such as humans is often challenged by the fact that teleosts experienced a genome duplication event in their early evolutionary history. Recent genome sequencing of three holosteans, the sister lineage to teleosts that did not experience a genome duplication event, has revealed these taxa to be critical for linking homologs between teleosts and other vertebrates. Sequencing of holostean genomes remains limited, thereby impeding further comparative genomic studies. Here we fill this sampling gap through the genomic sequencing of the longnose gar (*Lepisosteus osseus*). This annotated reference genome will provide a useful resource for a range of comparative genomic applications that span fields as diverse as immunogenetics, developmental biology, and the understanding of regulatory sequence evolution.

## Introduction

Teleost fishes represent over half of all living vertebrates and have successfully radiated in nearly all of the planet’s aquatic habitats (Ghezelayagh et al. 2022; Near et al. 2013). Teleosts are of vital ecological importance, form the basis of several multi-billion dollar industries (Lamberth & Turpie 2003; Sumaila et al. 2012; Mitcheson et al. 2013), and act as important model species (e.g. zebrafish and medaka) that are of high utility for human health research (Phillips & Westerfield 2014; Faillaci et al. 2018). The rapid accumulation of hundreds of genome sequences spanning the teleost Tree of Life has empowered unprecedented insights into the genomic basis for their evolutionary success (Daane et al. 2019), and provided key insights into teleost molecular biology with translational relevance to human health (Idilli et al. 2017). However, the development of both a deeper understanding of teleost genome evolution and the connection between teleost and human genomes has been challenged by the teleost-specific genome duplication (TGD) event that occurred during the early evolution of teleosts. This duplication event has complicated investigations of genomic novelty, homology, and synteny (Braasch et al. 2016; Hoegg et al. 2004). In contrast, the few living species of holosteans, non-teleost fishes (bowfin and gar) dubbed “living fossils” by Darwin (Darwin 1871) diverged from teleosts prior to the TGD (Thompson et al. 2021). Holostean genomes have been demonstrated to be critical for understanding gene synteny and homology of complex genomic regions between teleosts and other vertebrates (Thompson et al. 2021; Bi et al. 2021; Braasch et al. 2016). Being the closest living relatives of teleosts, holosteans provide particularly informative context for understanding whether genomic novelties identified in teleosts are in fact unique to teleosts and for linking teleost and other vertebrate genomes (Dornburg & Yoder 2022; Dornburg et al. 2021; Thompson et al. 2021; Braasch et al. 2016).

Extant holosteans include seven species of Lepisosteidae (gar; (Eschmeyer 1998)) and two species of Amiidae (bowfin; (Brownstein et al. 2022; Wright et al. 2022)). Analyses of the spotted gar (*Lepisosteus oculatus*) genome demonstrated the potential of holostean genomes for comparative studies, providing critical insights into the evolution of vertebrate immunity, development, and the function of regulatory sequences (Braasch et al. 2016). Recently, the alligator gar (*Atractosteus spatula*) genome was incorporated into an analysis of how vertebrates made the transition from water to land (Bi et al. 2021), and the genome of the distantly related eyetail bowfin (*Amia ocellicauda*, previously *Amia calva*, see (Brownstein et al. 2022)), provided understanding into other aspects of early vertebrate diversification including the evolution of scales, loci associated with the vertebrate adaptive immune response, and numerous other traits (Thompson et al. 2021). These studies have been imperative for our understanding of vertebrate evolution and molecular biology. However, they also underscore the potential insights that genomic sequencing of the remaining holostean genomes would provide. In particular, sequencing additional holostean species, with more focused investigations of within-clade sequence evolution, would facilitate a better understanding of highly fragmented regions in currently available holostean genome assemblies. In this study we present a high-quality assembly and annotation for the longnose gar (*Lepisosteus osseus*). This fourth holostean genome fills a critical sampling gap among holosteans, providing a valuable resource for genomic investigations of early vertebrate evolution as well as the necessary context for bridging research between model teleosts and the human genome.

## Results and Discussion

### Assembly and coverage of universal orthologs

Here we report a high-quality assembly of the longnose gar (**Figure 1A**) genome (**Supplemental Table S1** and NCBI Bioproject PRJNA811181). Dovetail Genomics (Scotts Valley, CA) performed DNA extraction from a longnose gar blood sample, library preparation, sequencing, and genome assembly. Genomic DNA was extracted using a Qiagen Blood and Cell Culture DNA Midi Kit (Germantown, MD), yielding DNA with an average fragment length of 95 kbp that was used in the construction of Chicago HiC sequencing libraries. The 10X supernova assembly resulted in 27,738 scaffolds forming a total final genome size of 1,014,182,714 bp, with 2.4% of the genome (24,076,280 bp) comprised of the ambiguous base ‘N’ and a GC content of 40.1%. During Dovetail Hi-Rise assembly the input assembly was further incorporated into 22,745 longer scaffolds. The total length of the resulting Dovetail Hi-Rise assembly was 1014.9 Mbp, with a contig N_50_ of 116.6 kbp. The N_50_ of the assembly was 53.0 Mbp scaffolds with a L_50_ of 8 scaffolds. This is similar to the spotted gar genome that is 945 Mbp long, with a contig N_50_ size of 68.3 kbp, and a scaffold N_50_ size of 6.9 Mbp (Braasch et al. 2016) and the eyetail bowfin reference genome which is 527 Mbp, with a scaffold N_50_ of 41.2 Mbp, an L_50_ of 9 scaffolds, and contig N_50_ of 21.1 kbp (Thompson et al. 2021) (**Figure 1B**). The L_90_ based on this assembly is 26 which is comparable to the spotted gar, where a chromosomal spread suggested 29 linkage groups (Braasch et al. 2016).

**Figure 1.**
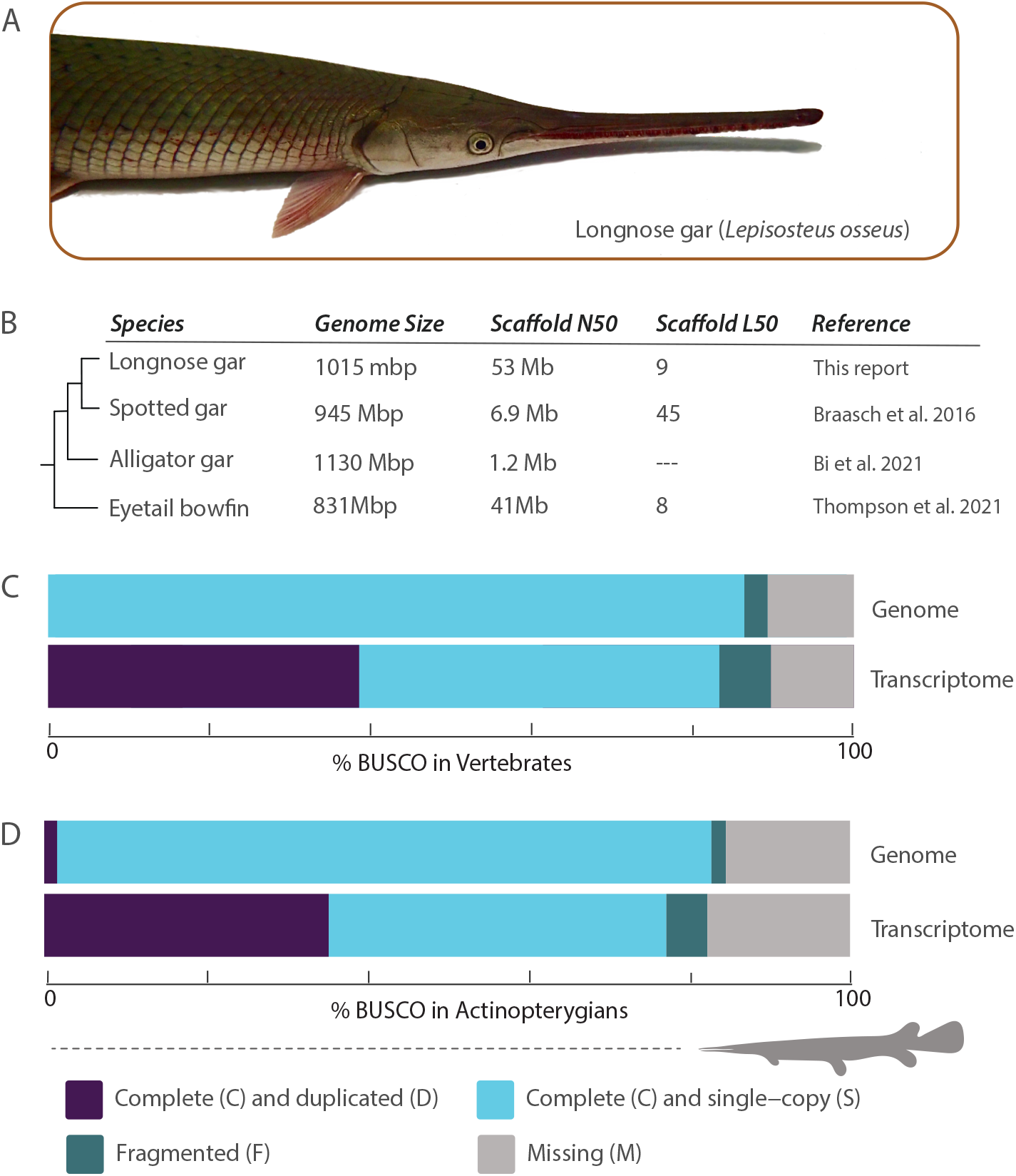
The annotated longnose gar genome.(A) Photo of a longnose gar wild-caught from North Carolina. (B) Comparison of holostean genome metrics. (C) BUSCO scores of the longnose gar genome and transcriptome compared to vertebrates (D) and actinopterygians. Photo credit: AD.

Benchmarking Universal Single-Copy Orthologs (BUSCO) analysis (Manni et al. 2021) comparing the longnose gar genome against the Actinopterygii dataset recovered a total of 3082 out of 3640 loci. Of these, 3017 (82.8%) are complete (2957 (81.2%) complete and single copy, 60 (1.6%) complete and duplicated), 65 (1.8%) fragmented, and the remaining 558 (15.4%) missing (**Figure 1C** and **Supplemental Table S2**). These numbers change slightly when compared to the Vertebrata BUSCO dataset. Out of 3354 total BUSCO groups, we recovered 2898 (86.4%) complete sequences [(2867 (85.5%) complete and single copy, 30 (0.9%) complete and duplicated], 97 (2.9%) fragmented, and 359 (10.7%) missing (**Figure 1D** and **Supplemental Table S2**). The higher number of sequences recovered when using the vertebrate versus actinopterygian databases mirrors a similar near twofold difference in missing data in the bowfin genome (Thompson et al. 2021). This may reflect a teleost bias for Actinopteryigan BUSCOs, or stem from a difference in the number of target loci. More work is needed to assess if the development of a BUSCO dataset for early diverging, non-teleost, actinopterygians is warranted.

### New insights into the transposable elements of Holostean genomes

Our RepeatModeler analysis (Flynn et al. 2020) reveals 29.1% of the longnose gar genome is composed of transposable elements (TEs; **Figure 2A**). Retroelements account for 14.02% of the transposons, 2.1 % of which are short interspersed nuclear elements (SINEs) and 6.5% of which are long interspersed nuclear elements (LINEs). Among LINEs, L2/CR1/Rex elements represent 4.3% of the total diversity, while L1/CIN4 elements represent only 0.4% (**Figure 2A**). DNA transposons cover 3.7% of the genome, with Tc1-IS630-Pogo elements reflecting 2.7% of the total diversity. In general SINES and LINES are some of the most frequent TE types (**Figure 2B**). Our results are on par with the 20% TE content found in the spotted gar, and contrasts with the levels of near 50% TE content in humans or zebrafish (*Danio rerio)* (**Figure 2C**) (Braasch et al. 2016). Similar to spotted gar, we find a high diversity of eukaryote TEs in the longnose gar genome after conducting a BLAST search against the Repbase (Jurka et al. 2005) database. 235 sequences of repeats matched to Repbase: 46 DNA transposons, 71 LINEs, 25 SINEs, 28 Long Terminal Repeats (LTRs), 20 tRNA and 37 sequences classified as Unknown. Our finding of CR1 parallels a similar discovery in the spotted gar, which was used as evidence to suggest that the absence of CR1 is a teleost-specific loss, and not a general condition of early ray-finned fish (**Figure 2A**) (Braasch et al. 2016). Additionally, our finding of CR1, L2, Rex1, and Babar reveals longnose gar to be the second known vertebrate with all four CR1-like families. As the only other known vertebrate with all four CR1-like families is the spotted gar, this finding highlights the potential utility of holosteans for understanding the evolution of early vertebrate TEs.

**Figure 2.**
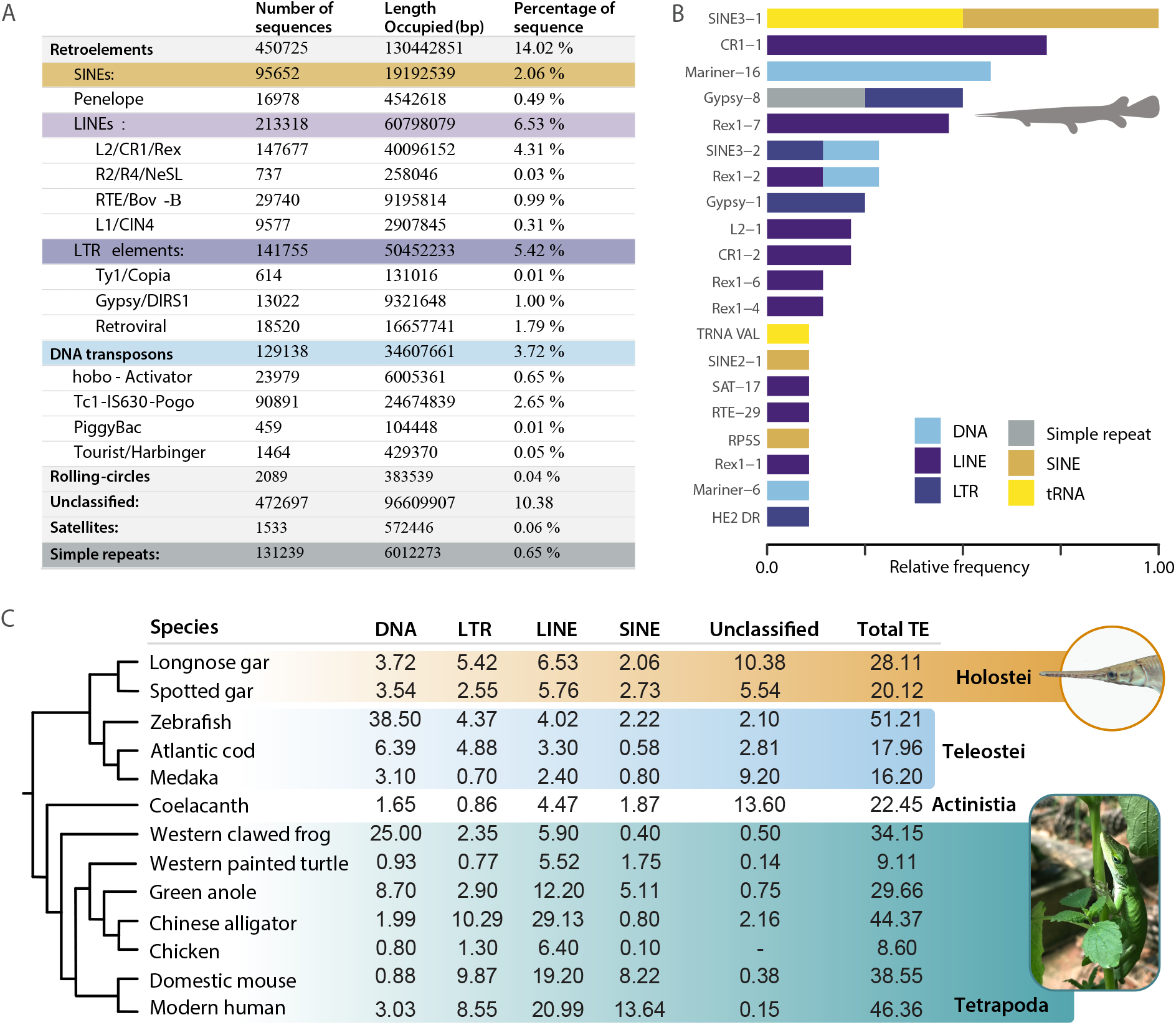
Overview of repetitive sequences within the longnose gar genome. (A) The distribution of repeat types identified from the longnose gar genome using RepeatModeler (Flynn et al. 2020). (B) The relative frequency of the 20 most frequent TEs that matched to Repbase. Colors correspond to TE types. (C) Comparison of TE class (DNA, LTR, LINE and SINES) percentages in the longnose gar with other vertebrates. Values for other vertebrates were obtained from literature: spotted gar (Braasch et al. 2016); zebrafish (Howe et al. 2013); Atlantic cod (Star et al. 2011); medaka (Kasahara et al. 2007); coelacanth (Amemiya et al. 2013); Western clawed frog (Hellsten et al. 2010); western painted turtle (Shaffer et al. 2013); green anole (Alföldi et al. 2011); Chinese alligator (Wan et al. 2013); Chicken (International Chicken Genome Sequencing Consortium 2004); domestic mouse (Mouse Genome Sequencing Consortium et al. 2002); and modern human (Lander et al. 2001). Photo credits: AD.

### Transcriptome sequencing and gene ontology

Transcriptome sequences were assembled from RNA-Seq derived from immune tissues of the same longnose gar individual from which the genome originated. The transcriptome sequences were *de novo* assembled using Trinity (Grabherr et al. 2011; Haas et al. 2013) and mapped to the genome using HISAT2. Processed reads were mapped to the genome and assessed using samtools v.1.18, with 30901143 out of 44987853 reads (68.7%) mapping to the genome. BUSCO (Manni et al. 2021) analysis was employed on the assembled transcriptomic sequences to quantify the completeness of the transcriptome (**Figure 1** and **Supplemental Table S3**). Comparing the longnose gar transcriptome to the Actinopterygii and Vertebrata databases yielded 2809 (77.2%) and 2792 (83.3%) complete sequences, respectively. The vertebrate database yielded a higher number of complete and single-copy (44.7%) and complete and duplicated (38.6%) sequences than the comparison to the actinopterygian database (41.9% and 35.3% respectively).

The transcriptome was used to annotate the genome (Zenodo DOI: <u>10.5281/zenodo.7435126</u>), predicting 30,068 proteins. This is similar to the 25,645 proteins predicted in the spotted gar genome by MAKER (Braasch et al. 2016). Gene ontology analysis of transcripts from pooled spleen, kidney and intestine RNA reveals the top *molecular functions* to include cytokine activity, signaling receptor regulator activity, signaling receptor activator activity and also include proteasomal complexes in the *cellular components*. These are all reflective of the immunological roles of these tissues (see **Supplemental Figures 1-3)**.

## Methods and Materials

### Sample acquisition

All research involving live animals was performed in accordance with relevant institutional and national guidelines and regulations, and was approved by the North Carolina State University Institutional Animal Care and Use Committee. The longnose gar specimen was wild-caught on the Haw River, North Carolina (35.626174, -79.057769) by the NC Wildlife Resources Commission using standard boat electroshocking methods and housed at the NC State University College of Veterinary Medicine in a 300 gallon tank with recirculating water at 18-23 °C. The individual was anesthetized using MS-222 and 2.5 mL of blood was collected into 0.5 mL of 87 mM EDTA for genomic sequencing. The fish was euthanized, and supplemental tissue samples (spleen, kidney and intestine) were collected for transcriptome sequencing.

### Chicago and Dovetail Hi-C library prep and sequencing

A Chicago library was prepared by Dovetail Genomics using ∼500 ng genomic DNA and methods described in (Putnam et al. 2016). In brief, chromatin was reconstituted *in vitro* and crosslinked with formaldehyde. Chromatin was digested (*Dpn*II) and the resulting 5’ overhangs were filled with biotinylated nucleotides. Blunt ends were ligated and DNA was purified. Purified DNA was obtained from protein by reversing the crosslinks and subsequently treated to remove the biotin that was not initially internal to the ligated fragments. The Hi-C library was then created using the methods as described in (Lieberman-Aiden et al. 2009) shearing the DNA to ∼350 bp mean fragment size. The sequencing libraries were generated using NEBNext Ultra enzymes and Illumina compatible adapters. Biotin-containing fragments were isolated using streptavidin beads before PCR enrichment of each library. The libraries were sequenced on an Illumina HiSeqX and yielded 163 million paired end reads (2 × 150 bp) that provided 7514.8X physical coverage of the genome (10-10,000 kbp).

### Scaffolding the assembly with Hi-Rise

Dovetail staff used HiRise (Putnam et al. 2016) to scaffold genome assemblies. The *de novo* assembly, Chicago library reads and the Dovetail HiC library reads were used as inputs for HiRise. The shotgun and Chicago library sequences were first aligned to the draft input assembly using a modified SNAP read mapper (http://snap.cs.berkeley.edu), that masked base pairs that followed a restriction enzyme junction. The Chicago data was aligned and scaffolded following aligning and scaffolding of the dovetail HiC library. Once all the sequences were aligned and scaffolded, shotgun sequences were used to close the gaps between contigs.

### Contamination removal and species verification

Contaminated and adaptor sequences were identified with feedback from NCBI and removed using custom scripts. The species’ identity was confirmed using tBLASTn searches with the universal barcode for fish species Cytochrome c Oxidase I (COI) as a query (Ivanova et al. 2007). Custom scripts have been archived on Zenodo (DOI: <u>10.5281/zenodo.7435126)</u>

### RNA sequencing

RNA was extracted (Qiagen RNeasy kit) from the spleen, kidney and intestine of the same individual longnose gar as the genome sequence and quantity and integrity of the isolated RNA were assessed using a NanoDrop 1000 (Thermo Fisher) and Agilent Bioanalyzer respectively. In brief, mRNA was enriched using oligo(dT) beads, rRNA was removed using a Ribo-Zero kit (Epicentre, Madison, WI) and mRNA was randomly fragmented. Each RNA extraction was equalized for a final concentration of 180 ng/µL. Library preparation and sequencing was performed by Novogen Corporation (Sacramento, CA). Next-gen sequencing (2 × 150 bp paired end reads) was performed on a NovaSeq 6000 instrument (Illumina). Adapter sequences and poor quality reads were filtered with Trimmomatic v34 (Bolger et al. 2014). The transcriptome was *de novo* assembled with Trinity v2.14.0 (Grabherr et al. 2011). Completeness of the transcriptomes was assessed using a BUSCO analysis (Manni et al. 2021). Raw reads and computationally assembled transcriptome sequences were deposited onto NCBI under the accession numbers SRR19528583 and GKEG00000000, respectively.

Transcriptome sequences were further analyzed to assign gene ontology. The assembled RNA-seq from Trinity was translated using Transdecoder to identify the candidate coding regions in the transcript sequences. The longest open reading frames (orfs) output was used for BLASTx and BLASTp analysis against the uniprot database (Nov 2021 release) to get the top target sequences for every transcript. Hmmscan v.3.3.2 was used to search for protein sequences in the Pfam-A (Nov 2021 release) library. Signalp v.5.0b (Teufel et al. 2022) and TMHMM v.2.0c (Krogh et al. 2001) were used to identify the signal peptides and the transmembrane proteins. Trinotate v.3.2.2 used sqlite database along with the Trinity assembled transcriptome and the longest orfs from Transdecoder to create a gene transmap (Bryant et al. 2017). The transcripts were finally annotated using Trinotate and then further analyzed to obtain the GO annotations. The GO terms were visualized using the enrichplot and ggupset packages in R. All code has been made available on Zenodo: (DOI: <u>10.5281/zenodo.7435126</u>).

### Annotation and genome quality assessments

BRAKER2 (Brůna et al. 2021) was used to annotate the longnose gar genome, which uses GeneMark-ET (Lomsadze et al. 2014) to predict the preliminary genes and generate a genemark.gtf output that was used for training with Augustus (Stanke et al. 2006). The genome was filtered to remove any duplicates and adapters. The transcriptome sequences were aligned using HISAT2 (Kim et al. 2019) to get an aligned sorted bam file. RepeatModeler (Flynn et al. 2020) identified the repeats in the genome to prevent mis-annotation of the repeats as protein coding genes. The consensus file containing repeats was used as input for RepeatMasker (Chen 2004) to soft-mask the repeats for BRAKER. The masked genome and the aligned transcriptomes were loaded into BRAKER to obtain the annotated proteins.

## Supplemental Information

Supplemental figures and tables are available in the pdf file included with this manuscript.

## Data availability

The longnose gar genome sequence is available through NCBI Bioproject PRJNA811181. Raw transcriptome reads and computationally assembled transcriptome sequences are available through NCBI under the accession numbers SRR19528583 and GKEG00000000, respectively. All code used for analyses and the BRAKER2 genome annotation are available on Zenodo (DOI: <u>10.5281/zenodo.7435126</u>)

## Acknowledgements

We thank Ingo Braasch and Andrew Thompson (Michigan State University) for very helpful discussions on genome analyses strategies, Kent Passingham (NC State University) for assistance with blood collection, and the NCBI staff for assistance with identifying adapter and other contaminating sequences. We thank A. Bogan for facilitating the photography of the *Anolis* in Figure 2.

## Author contribution

JAY and AD conceived of the project; MF, DJW, JAY and AD collected *Lepisosteus osseus*; RM executed genomic analyses including analysis of repetitive elements and genome annotation; DJW and RM assembled transcriptome sequences; RM, KBC and AD analyzed transcriptome sequences; RM conducted transcriptome annotation; RM and AD visualized data; RM, KBC, JAY and AD wrote the first draft of the manuscript; all other authors contributed to the subsequent writing; JAY and AD supervised the project. All authors read and approved the manuscript.

## Funding

This work was supported by the National Science Foundation (IOS1755330 to JAY and IOS1755242 to AD), the National Evolutionary Synthesis Center, NSF EF0905606 (DJW), and funding from the Triangle Center for Evolutionary Medicine (AD and JAY).

## Notes

### Competing Interest Statement

The authors have declared no competing interest.

