## Supplemental Information for "A chromosome level genome assembly of longnose gar, *Lepisosteus osseus*"

*Genome Biology and Evolution*

### Contents:

Supplemental Table S1. Genome assembly statistics of longnose gar. 2

Supplemental Table S2. BUSCO analysis of the longnose gar genome assembly. 2

Supplemental Table S3. BUSCO analysis of the longnose gar transcriptome assembly. 3

Supplemental Figure S1. Summary of Gene Ontology Molecular Functions analysis from the longnose gar transcriptome. 4

Supplemental Figure S2. Summary of Gene Ontology Biological Processes analysis from the longnose gar transcriptome. 5

Supplemental Figure S3. Summary of Gene Ontology Cellular Components analysis from the longnose gar transcriptome. 6

**Supplemental Table S1.** Genome assembly statistics of longnose gar

| Feature | Value |
|---|---|
| GC content | 40.1% |
| Number of Scaffolds | 22,745 |
| Number of scaffolds >1 kbp | 22,709 |
| Contig $N_{50}$ | 116.61 kb |
| Scaffold $N_{50}$ | 52.996Mb |
| Scaffold $L_{50}$ | 8 |
| $L_{90}$ | 26 scaffolds |
| $N_{90}$ | 5.560 Mb |
| Longest scaffold | 74,198,471 bp |
| Number of gaps | 27,358 |
| Percent of genome in gaps | 2.45% |

**Supplemental Table S2.** BUSCO analysis of the longnose gar genome assembly

| Feature | Actinopterygii | Vertebrata |
|---|---|---|
| Complete BUSCOs (C) | 3017 (82.8%) | 2898 (86.4%) |
| Complete and single-copy BUSCOs (S) | 2957 (81.2%) | 2867 (85.5%) |
| Complete and duplicated BUSCOs (D) | 60 (1.6%) | 30 (0.9%) |
| Fragmented BUSCOs (F) | 65 (1.8%) | 97 (2.9%) |
| Missing BUSCOs (M) | 558 (15.4%) | 359 (10.7%) |
| Total BUSCO groups searched (n) | 3640 | 3354 |

**Supplemental Table S3.** BUSCO analysis of the longnose gar transcriptome assembly

| Feature | Actinopterygii | Vertebrata |
|---|---|---|
| Complete BUSCOs (C) | 2809 (77.2%) | 2792 (83.3%) |
| Complete and single-copy BUSCOs (S) | 1524 (41.9%) | 1492 (44.7%) |
| Complete and duplicated BUSCOs (D) | 1285 (35.3%) | 1294 (38.6%) |
| Fragmented BUSCOs (F) | 187 (5.1%) | 214 (6.4%) |
| Missing BUSCOs (M) | 644 (17.7%) | 348 (10.3%) |
| Total BUSCO groups searched (n) | 3640 | 3354 |

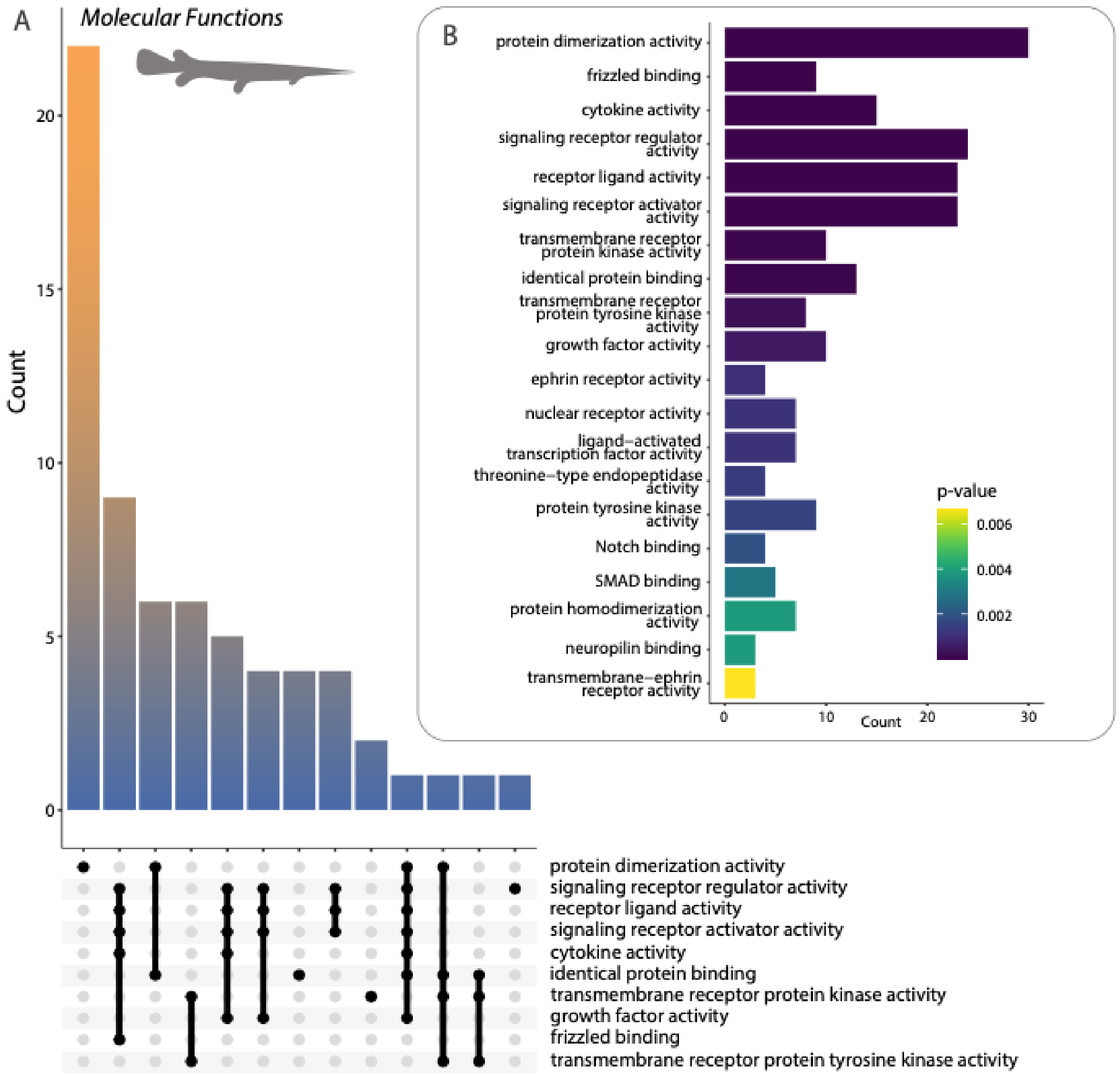


**Supplemental Figure S1.** Summary of Gene Ontology *Molecular Functions* analysis from the longnose gar transcriptome.
Predicted proteins from the transcriptome were used as inputs to assess (A) functions and their intersections and (B) most common terms.

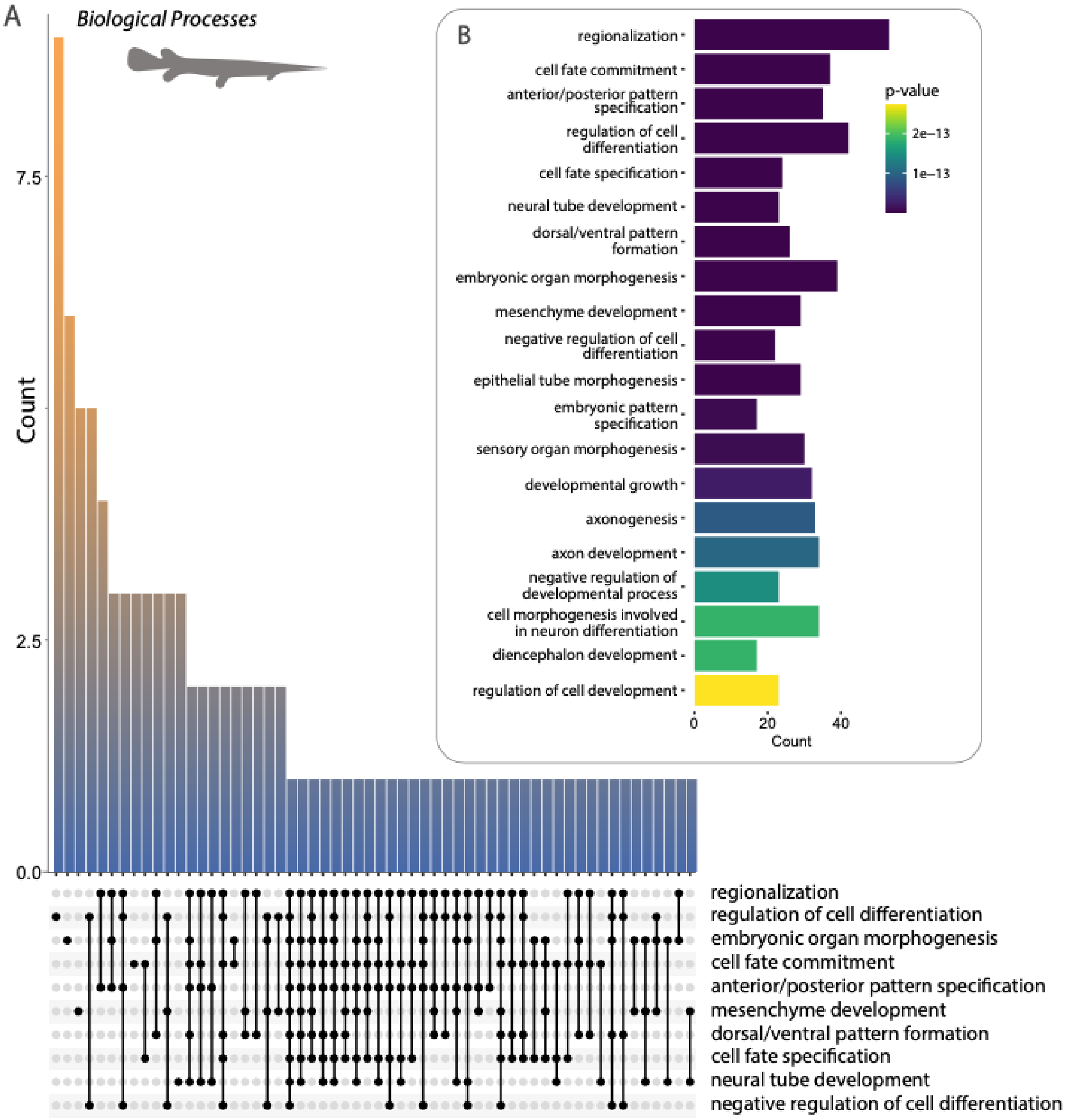


**Supplemental Figure S2.** Summary of Gene Ontology *Biological Processes* analysis from the longnose gar transcriptome.
Predicted proteins from the transcriptome were used as inputs to assess (A) processes and their intersections and (B) most common terms.

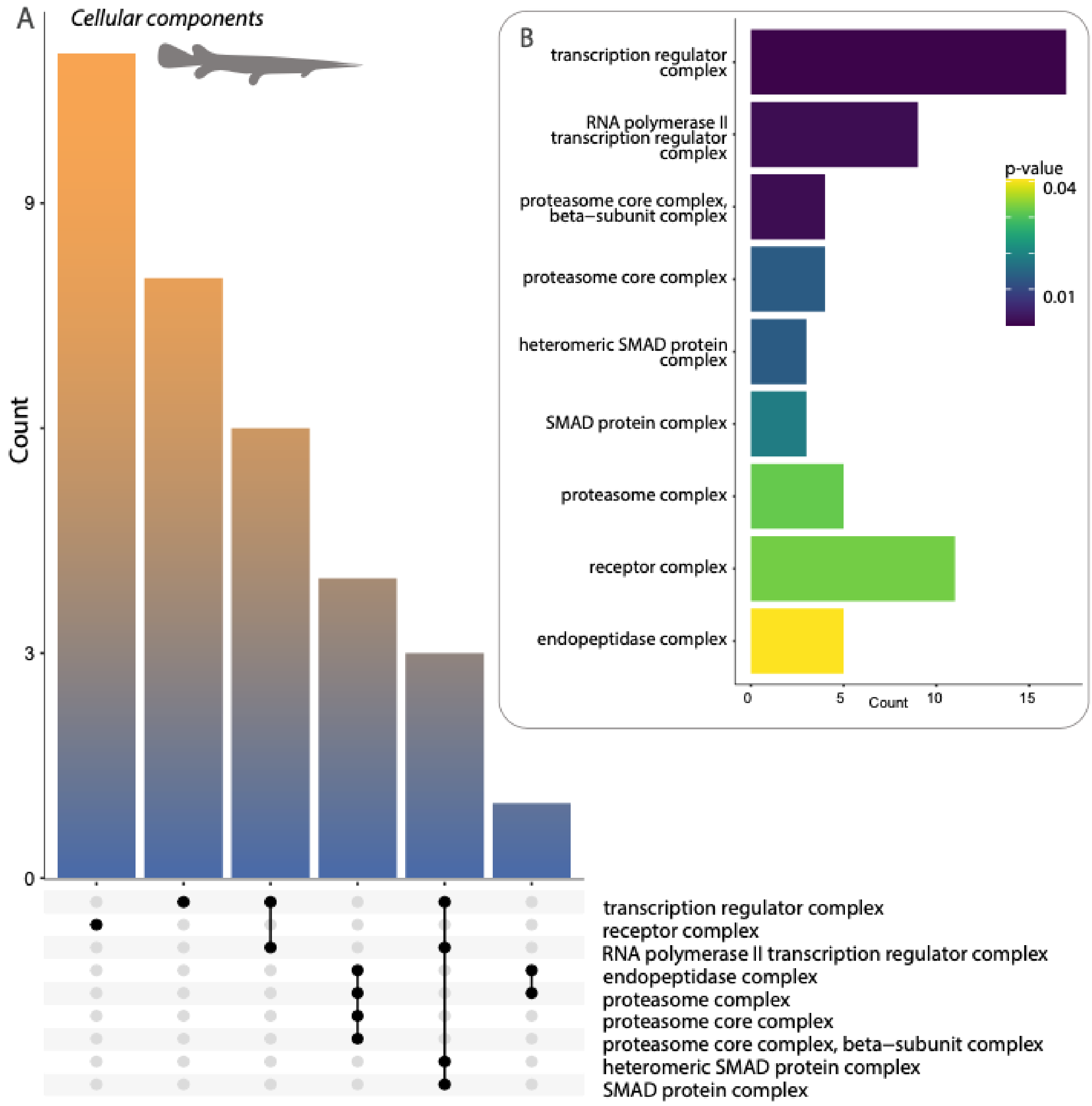


**Supplemental Figure S3.** Summary of Gene Ontology *Cellular Components* analysis from the longnose gar transcriptome.
Predicted proteins from the transcriptome were used as inputs to assess (A) components and their intersections and (B) most common terms.
